# Grapevine virus L: a Novel Vitivirus in Grapevine

**DOI:** 10.1101/314674

**Authors:** Humberto Debat, Diego Zavallo, Reid Soltero Brisbane, Darko Vončina, Rodrigo P.P. Almeida, Arnaud G. Blouin, Maher Al Rwahnih, Sebastian Gomez-Talquenca, Sebastian Asurmendi

**Affiliations:** Instituto de Patología Vegetal, Centro de Investigaciones Agropecuarias, Instituto Nacional de Tecnología Agropecuaria (IPAVE-CIAP-INTA), Córdoba, Argentina, X5020ICA; Instituto de Biotecnología, Centro de Investigación en Ciencias Veterinarias y Agronómicas (IB-CICVyAINTA), Buenos Aires, Argentina, 1686; Foundation Plant Services, Davis, CA 95616, USA; Department of Plant Pathology, Faculty of Agriculture, University of Zagreb, Zagreb, Croatia; Department of Environmental Science, Policy and Management, University of California, Berkeley, CA, USA; The New Zealand Institute for Plant & Food Research Limited, Private Bag 92169, Auckland, 1142, New Zealand; Department of Plant Pathology, University of California, Davis, CA, USA; Estación Experimental Agropecuaria Mendoza, Instituto Nacional de Tecnología Agropecuaria (EEAMendoza-INTA), Luján de Cuyo, Mendoza, Argentina, 5534; CONICET, Argentina

**Author notes:** These authors contributed equally to this work (Humberto Debat, Diego Zavallo). Correspondence: Humberto Debat, Sebastian Gomez-Talquenca. **Accession numbers**: Grapevine virus L sequences have been deposited in GenBank under accession numbers: MH248020 (GVL-RI), MH643739 (GVL-KA), MH681991 (GVL-VL), MH686191 (GVL-SB).

**Keywords:** *Vitivirus*, Grapevine, virus discovery, *Betaflexiviridae*

## Abstract

*Vitivirus* are ssRNA(+) viruses in the family *Betaflexiviridae* (subfamily *Trivirinae*). There are currently ten ICTV recognized virus species in the genus; nevertheless, the extended use of NGS technologies is rapidly expanding their diversity and six more have been proposed recently. Here, we present the characterization of a novel virus from grapevines, which fits the genomic architecture and evolutionary constraints to be classifiable within the *Vitivirus* genus. The detected virus sequence is 7,607 nt long, including a typical genome organization of ORFs encoding a replicase (RP), a 22 kDa protein, a movement protein, a coat protein (CP) and a nucleic acid binding protein. Here, we present the characterization of a novel virus from grapevines. Phylogenetic analyses based on the predicted RP and CP protein unequivocally places the new virus within the *Vitivirus* genus. Multiple independent RNAseq data confirmed the presence of the detected virus in berries at diverse developmental stages. Additionally, we detected, confirmed, and assembled virus sequences from grapevine samples of distinct cultivars from America, Europe, Asia and Oceania, sharing 74.9%-97.9% nt identity, suggesting that the identified virus is widely distributed and diverse. We propose the name grapevine virus L (GVL) to the detected *Vitivirus*.

## Introduction

Vitiviruses have flexuous, non-enveloped, filamentous virus particles of 725-785 nm and 12 nm in length and diameter, respectively, with a nucleocapsid that is cross-banded and diagonally striated. Vitiviruses have a linear ssRNA(+) genome (∼7.3-7.6 kb), with a methylated nucleotide cap at the 5’ end and a 3’ poly (A) tail [1-2]. There are ten species of vitiviruses recognized by the ICTV, nevertheless, six new species have been proposed recently, five of those infecting grapevine (*Vitis vinifera*) [3-7]. Grapevine is the most prevalent natural host of vitiviruses, but they have also been found to infect several crops such as mint (*Mentha x glaciaris*), arracacha (*Arracacia xanthorrhiza*) blue agave (*Agave tequilana*), kiwi (*Actinidia chinense*) and blackberry (Rubus spp) [8-11]. Vitiviruses appear to be latent in *V. vinìfera* cultivars, and so far, only *Grapevine virus A* and *Grapevine virus B* have been consistently associated to grapevine diseases of the rugose wood complex (Grapevine vitiviruses) or Shiraz disease (reviewed by Minafra [12]). The etiological role of the recently described vitiviruses should be assessed in order to establish its relationships with known or unknown viral diseases. In addition, the synergistic effects between vitiviruses and other grapevine viruses appears to be significant [13]. The availability of complete sequences of these viruses could allow the development of full-length infectious clones, fulfillment of the Koch’s postulates, and improve our understanding of the biological role of these viruses [14]. Here, we present the characterization of a novel vitivirus for which we tentatively propose as grapevine virus L (GVL). We were able to detect and characterize GVL in several distinctive *Vitis vinifera* cultivars from four continents.

## Material and Methods

Virus discovery, confirmation, and annotation were implemented as described in [4-5, 15-16]. RT-PCR assays were deployed as reported in Al Rwahnih et al. [17]. Raw RNA data from several Sequence Read Archive (SRA) accessions was downloaded from the National Center for Biotechnology Information database (NCBI). These files was processed in the following form: trimmed and filtered with the Trimmomatic tool as implemented in http://www.usadellab.org/cms/?page=trimmomatic, and the resulting reads of each library were assembled *de novo* with Trinity v2.6.6 release with standard parameters [18]. The obtained transcripts were subjected to bulk local blastx searches (E-value < 1e-5) against a refseq virus protein database available at ftp://ftp.ncbi.nlm.nih.gov/refseq/release/viral/viral.1.protein.faa.gz. The resulting hits were explored manually and curated by iterative mapping using Bowtie2 available at http://bowtie-bio.sourceforge.net/bowtie2/. ORFs were predicted by ORFfinder (https://www.ncbi.nlm.nih.gov/orffinder/) and annotated with the NCBI conserved domain search tool as implemented in https://www.ncbi.nlm.nih.gov/Structure/cdd. Phylogenetic insights were generated by multiple amino acid alignments of replicase proteins (BLOSUM62 scoring matrix) using as best-fit algorithm E-INS-i, which drives local alignment with generalized affine gap costs, and thus is applicable for RNA polymerases where diverse domains are dispersed among several highly divergent regions, by MAFTT v7.394 as implemented in https://mafft.cbrc.jp/alignment/software/. The aligned proteins were subsequently used as input for FastTree 2.1.5 http://www.microbesonline.org/fasttree/maximum likelihood phylogenetic trees (best-fit model = JTT-Jones-Taylor-Thorton with single rate of evolution for each site = CAT) computing local support values with the Shimodaira-Hasegawa test (SH) and 1,000 resamples. SNPs were detected by the FreeBayes v0.9.18S tool with standard parameters implemented in https://github.com/ekg/freebayes.

## Results and Discussion

In order to explore the potential grapevine viral landscape on High Throughput Sequencing (HTS) publicly available libraries, we downloaded raw RNA data from several SRA accessions on the NCBI database, finding known virus in the vast majority of tested samples. The raw data were assembled *de novo* and the obtained transcripts were subjected to bulk local blastx searches (E-value < 1e-5) against a refseq virus protein database. The resulting hits were explored manually. One dataset, specifically NCBI Bioproject PRJNA400621, SRX3144921-SRX3144956, composed of RNA libraries from grape berry samples of *V. vinifera* cv. Riesling and cv. Cabernet sauvignon from Beijing, China [19], presented a 6,155 nt long transcript from the assembled transcriptome of the SRX3144956 library. This sequence obtained a highly significant hit (E-value = 0; sequence identity 69%) with the replicase protein of *Grapevine virus E* (GVE, YP_002117775.1, [20]). Iterative read mapping allowed to extend and polish the assembled contig into a 7,607 nt long sequence, supported by 25,112 reads and a mean coverage of 161.7X, sharing the genome organization of vitiviruses (**Figure 1.A**; **Table 1**). The assembled virus sequence has a pairwise identity with GVE of 68.2% at the genomic RNA level. The predicted structure of the detected virus is conformed by a 67 nt 5′UTR, followed by five ORFs and a 66 nt 3′UTR, excluding the poly (A) tail of at least three adenosines, and unknown length. Both predicted UTRs are highly similar in length and size in relation to other *Vitivirus*, more precisely GVE and the proposed GVI and GVG (**Supp. Figure 1**). ORF1 (67-5,158 nt coordinates) encodes a replicase protein (RP) of 1,696 aa (192.1 kDa), sharing a 70.2% with GVE replicase (NC_011106). We employed the NCBI conserved domain search tool to annotate the RP, which presents a characteristic domain architecture of *Vitivirus*. The 5’ region of RP presents a viral methyltransferase domain (E-value = 1.76e-56; Pfam = pfam01660; 46-337 aa coordinates), which has been involved in mRNA capping, followed by a DEXDc type viral RNA helicase (E-value = 1.47e-12; Pfam = pfam01443; 750-1047 aa coordinates). Interestingly, an Alkylated DNA repair dioxygenase domain (AlkB), of the2OG-Fe(II) oxygenase superfamily, was identified within the helicase domain (E-value = 2.34e-06; COG = COG3145; 917-1032 aa coordinates) as previously described for an American isolate of GVE [21]. Finally, as expected, an RNA dependent RNA polymerase domain was found at the 3’ region of the RP (RdRP_2; E-value = 3.47e-27; Pfam = pfam00978; 1313-1637 aa coordinates).

**Figure 1.**
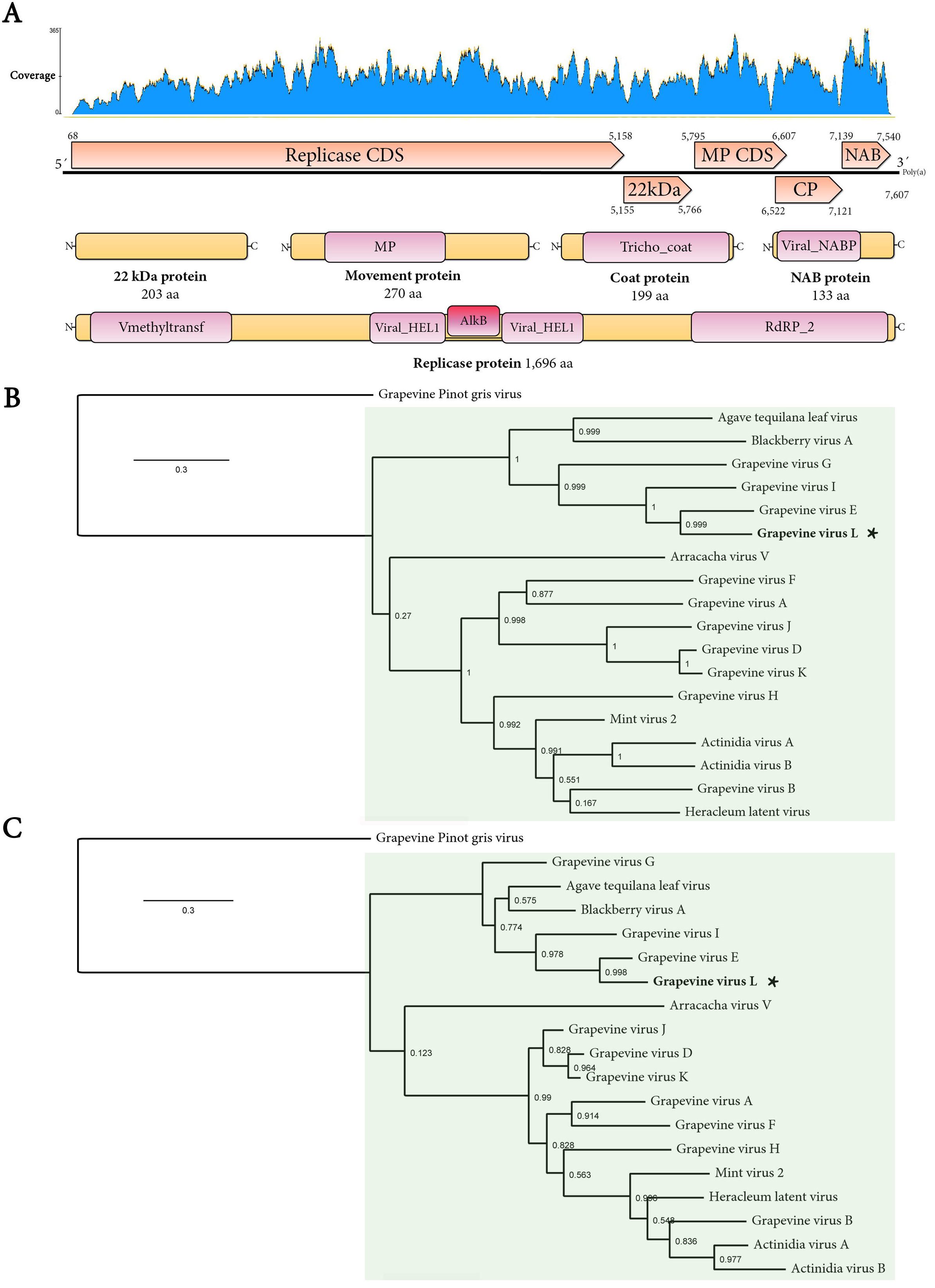
**(A)** Molecular characterization of grapevine virus L (GVL). Mapping pattern of virus RNA reads from library SRX3144956 to the consensus putative GVL assembly indicating coverage per position. The genome architecture of GVL is characterized by a monosegmented ssRNA(+) virus encoding for five ORFs arranged as 5’-UTR-RP-HP-MP-CP-NABP-UTR-3’. The predicted product of each ORF is depicted including the associated domains determined by NCBI-CDD. Abbreviations: RP, replicase protein; 22kDa, 22k Dalton hypothetical protein; MP, movement protein; CP, coat protein; NABP, nucleic acid binding protein. See the text for additional domain information. (**B**) Phylogenetic insights of putative grapevine virus L (GVL) in relation to accepted and proposed members of the genus *Vitivirus* based on MAFFT alignments (BLOSUM62 scoring matrix using as best-fit algorithm E-INS-i) of the predicted replicase and capsid protein (**C**) followed by maximum likelihood trees generated by FastTree (best-fit model = JTT-Jones-Taylor-Thorton with single rate of evolution for each site = CAT) computing local support values with the Shimodaira-Hasegawa test (SH) and 1,000 resamples). Accession numbers of the corresponding sequences: actinidia virus A (JN427014), actinidia virus B (NC_016404), mint virus 2 (AY913795), grapevine virus B (NC_003602), grapevine virus H (MF521889), grapevine virus F (NC_018458), grapevine virus A (NC_003604), grapevine virus K (NC_035202), arracacha virus V (NC_034264), grapevine virus E (NC_011106), agave tequilana leaf virus (NC_034833), grapevine virus D (KX828708), grapevine virus G (MF405923), grapevine virus I (MF927925), heracleum latent virus (X79270), and grapevine Pinot gris virus (*Trichovirus*, NC_015782) used as outgroup. Scale bar represents substitutions per site. Node labels are FastTree support values. Asterisk indicates GVL.

**Table 1.**
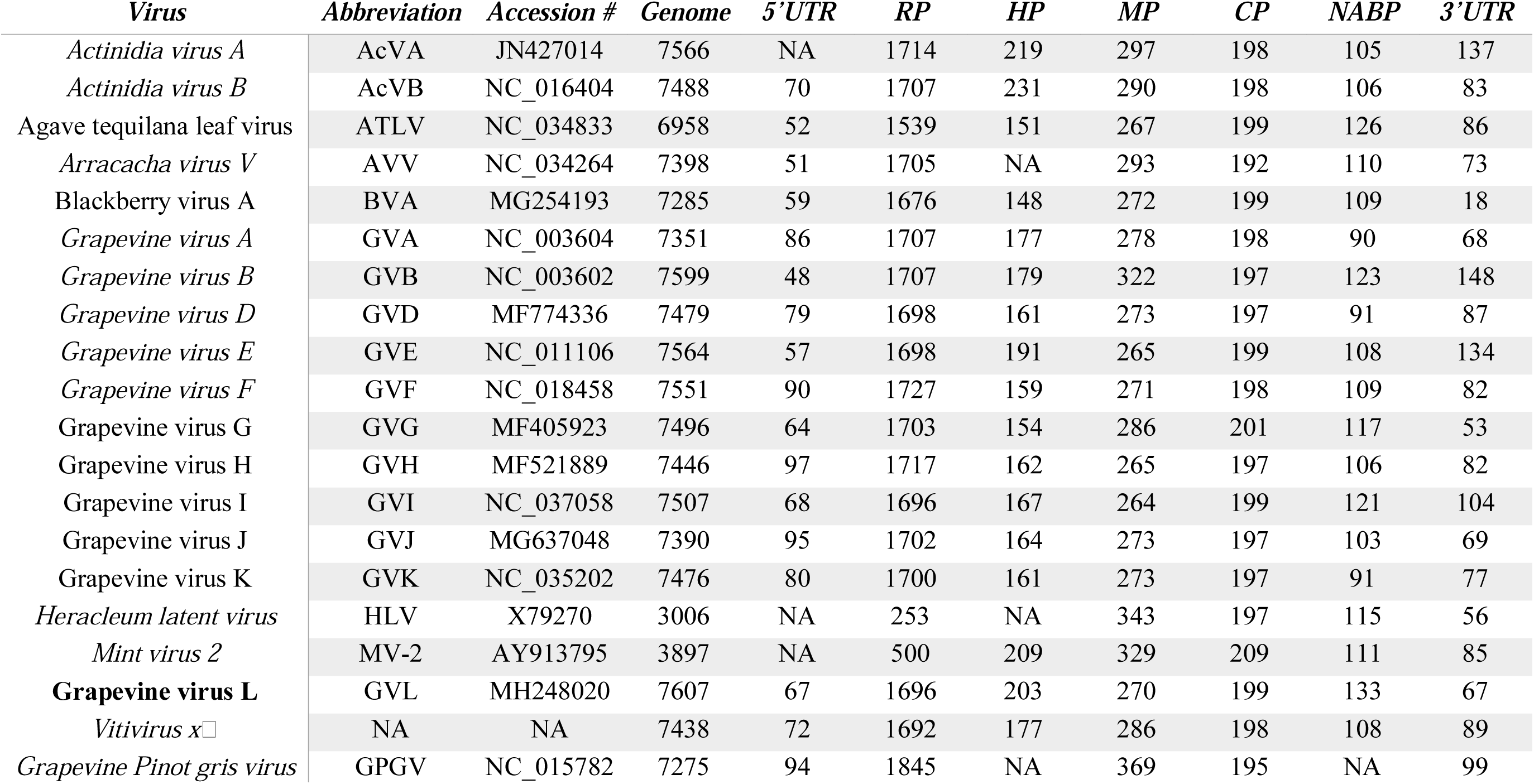
Diverse structural highlights of grapevine virus L in the context of proposed and ICTV recognized (in italics) *Vitivirus* species. *Grapevine Pinot gris virus* (genus *Trichovirus*) is included as outgroup for phylogenetic analyses. Accession #: NCBI Genbank accession number; Genome: Genome size (nt), 5′U: 5′UTR length (nt); RP: replicase protein length (aa); HP: hypothetical protein length (aa); MP: movement protein length (aa); CP: coat protein length (aa); NABP: nucleic acid binding protein length (aa); 3′UTR: 3′UTR length excluding the poly A tail (nt).

Although vitiviruses utilize a series of sub-genomic RNAs (sgRNA) as general strategy for expression of ORF2-5, it is worth mentioning that ORF2 (5,155-5,766 nt) of the detected virus overlaps in the tetranucleotide AUGA with ORF1. This overlapping cue might probably function as a termination–reinitiation strategy signal, which was determined for caliciviruses and victoriviruses for coupled start-stop translation [22]. This putative AUGA signal between ORF1-ORF2, although not mentioned, is present in the genome sequence of the recently proposed grapevine virus J [5]. The predicted ORF2 encodes a leucine rich (15.3% L) 22.8 kDa protein, of unknown function, which is the most highly divergent protein of the identified virus (44% similarity and 98% coverage to the ORF2 encoded hypothetical protein of GVE, AHB08905). ORF3 (5,795-6,607 nt) encodes a putative movement protein (MP) of 270 aa (29.7 kDa) with a viral movement protein domain (E-value = 6.61e-07; Pfam = pfam01107) between the 26-195 aa coordinates. ORF4 (6,522-7,121 nt), partially overlapping with ORF3, encodes a coat protein (CP) 199 aa long (21.4 kDa) presenting a *Trichovirus* coat protein-like domain (Tricho_coat; E-value = 4.26e-45; Pfam = pfam05892; 37-199 aa coordinates). The predicted CP of the identified virus shares 78.4% similarity with GVE-CP. Lastly, ORF5 (7,139-7,540 nt) encodes a 133 aa putative nucleic acid binding protein (15 kDa) with a nucleic acid binding domain (Viral_NABP; E-value = 1.40e-06; Pfam = pfam05515) between the 5-76 aa coordinates. This region shares significant similarity with the CTV_P23 domain (E-value =3.79e-06) of *Citrus tristeza virus* (*Closteroviridae*) P23 protein which is involved in asymmetrical RNA Accumulation [23].

The criteria for species demarcation in the family *Betaflexiviridae*, proposed by ICTV, is less than ca. 72% overall nucleotide identity and below 80% amino acid identity of CP or replicase predicted proteins [1]. Given that the detected virus shares 68.2% similarity with GVE at the genomic RNA level, and a 70.2% and 78.2% identity with the respective RP and CP proteins (**Supp. Figure 2**), is tentatively suggested that the detected virus sequence corresponds to a new species, for which we propose the name grapevine virus L (GVL). In order to entertain this hypothesis, based on genetic criteria, we gathered evolutionary insights of the putative GVL sequence identified in cv. Riesling grapevines from China (GVL-RI) in the context of the *Betaflexiviridae* family. We generated a phylogenetic tree based on RP proteins corresponding to the 93 refseq *Betaflexiviridae* RP sequences available at NCBI. Unequivocally, the putative GVL RP clustered within the *Vitivirus* genus (**Supp. Figure 3**). To provide local tree topology and additional phylogenetic insights we generated maximum likelihood phylogenetic trees of both the RP and CP protein of accepted and proposed members of the *Vitivirus* genus. In both cases, putative GVL clustered within one of the two major sub-clades, conformed by GVE and the proposed species GVG, GVI, and branching more distantly, the agave tequilana leaf virus (**Figure 1.B-C**). These results support the tentative assignment of the detected virus as a probable member of the *Vitivirus* genus.

To validate the putative GVL detection in SRX3144956 library, we took advantage of the data generated by Chen et al [19]. We mapped the 1,014,833,524 raw reads available at the PRJNA400621 project to the assembled GVL sequence, using as threshold two nt mismatches and a 22 nt seed and a cut-off value >10 reads hits per library with the Bowtie2 tool. None of the 18 RNA libraries of cv. Cabernet Sauvignon had the putative GVL. Interestingly, we were able to detect the virus in six additional cv Riesling libraries of the same study, obtained from independent biological samples conformed by grape berries sampled in three distinct developmental stages (E-L 35, E-L 36 and E-L 38) (**Supp. Table 1**). Moreover, while analyzing the intrinsic variability of the detected GVL sequences, we noticed that two of the samples presented 24 highly supported SNPs at frequencies >0.5 as detected by the FreeBayes v0.9.18S tool (**Supp. Table 2**). Given the low p-values of the variants and that 70% of the detected SNPs were silent, we speculate that these SNPs are not artifactual.

In order to explore the distribution of GVL, we gathered data from grapevine samples from Croatia [24], USA, and New Zealand. A collection of HTS libraries obtained from petioles RNA samples of autochthonous Croatian grapevines (cv Babica, cv Vlaska, cv Dobricic and cv Ljutun; Bioproject PRJNA415169) had virus RNA reads similar to GVL. In turn, *de novo* assembly and curation of the putative GVL transcript by iterative mapping resulted in a 7,584 nt virus genome (GVL-VL, mean coverage of 675X), sharing the genomic organization and a 75.1 % nt identity with GVL-RI (**Supp. Table 3**). The presence of this virus was subsequently confirmed by targeted RT-PCR (see below) of the HTS sequenced samples. Two PCR products were bi-directionally Sanger-sequenced and the obtained sequences shared a 100 % identity with GVL-VL.

The sample from USA corresponded to a quarantine selection Katelin (KA), which was received in 2012 for inclusion in the Foundation Plant Services (FPS, UC-Davis, CA) collection. The vine was grown in a screenhouse and assayed for known grapevine viruses as described previously [16]. In addition, total RNA from the positive sample was high-throughput sequenced and the obtained ca. 11.5 million raw reads were filtered and *de novo* assembled with parameters described in Al Rwahnih et al. 2018 [16]. A contig with high similarity with GVL was identified by BLAST and subsequently refined, resulting in a 7,591 nt virus draft genome (GVL-KA, MH643739), sharing the genomic architecture and a 97.9% nt identity with GVL-RI (**Supp. Table 3**).

Lastly, the presence of GVL was confirmed in an HTS library derived from total RNA isolated from cv. Sauvignon Blanc, which was originally collected in April 2016 in the region of Marlborough, South Island, New Zealand. The virus was detected in only one of the 225 Vitis sp. surveyed by HTS. The presence of the virus was confirmed by RT-PCR and Sanger sequencing. A draft genome with a partial truncation at 5’ was assembled, encompassing 7,365 nt, lacking both the 5’UTR and a ca. 184 nt coding region of the replicase (GVL-SB, mean coverage of 166X). The obtained sequence shared a 75.2% nt identity with GVL-RI. All draft genome sequences were annotated both structurally and functionally, showing consistent signatures of vitiviruses and a shared genome organization (**Supp. Figure 4**). In turn, sequence identity values were calculated indicating high sequence similarity in the predicted coat protein (92.5% to 99.5% aa identity) and significant divergence in the 22 kDa predicted protein (53.2% to 96.6% aa identity) (**Supp. Table 3**).

Further, an RT-PCR assay was deployed as reported in Al Rwahnih et al. [17] to investigate the prevalence of GVL in grapevine samples from USA, using forward [AGTTGAAGTCTAGGTGCACAC] and reverse [GTACTCAGACTTCCCCGATCTA] primers designed to target the MH248020 sequence of GVL-RI. The specificity of this primer pair in the detection of GVL was confirmed by RT-PCR using Total nucleic acid (TNA) from vines infected with GVA, GVB, GVD, GVE, GVH, GVG, GVJ. TNA was freshly extracted from leaf petiole tissue using the MagMax 96 Viral RNA isolation kit with the MagMax Express-96 magnetic particle processor (Thermo Fisher Scientific, USA). Using the above primers in those assays, none of the grapevines that were infected with the non-target vitiviruses were found to be positive, whereas a sample of grapevine cv. Katelin tested positive. The development of this specific RT-PCR assay will allow for the detection of GVL in field tests, and in clean-stock programs, facilitating a more effective control of this virus.

Given the intrinsic significant variability observed among the diverse GVL identified isolates, in the context of an overall interspecies affinity of GVL and GVE, we proceeded with additional phylogenetic analyses. We integrated the predicted products of four assembled GVL sequences and the five available, tentatively complete, genome sequences of GVE in multiple amino acid alignments (isolates TvAQ7, of *V. labruscana* from Japan; WAHH2, of *V. vinifera* cv Cabernet sauvignon from USA; SA94, of *V. vinifera* cv. Shiraz from South Africa; VVL-101, *V. vinifera* cv. Vlaska from Croatia and GFMG-1 from China). The maximum likelihood phylogenetic trees based both on RP and CP alignments mirrored the preceding tree topology presented in **Figure 1.B** and supported clade branching between GVL and GVE in the context of clustering within *Vitiviruses* (**Figure 2.A-B**). The availability of more RNA sequences corresponding to GVL would provide insights into the evolutionary trajectory and phylodynamics of GVL. The simultaneous detection of GVL in the Americas, Europe, Asia and Oceania in diverse *V. vinifera* cultivars suggest that this novel virus is widely distributed but not widely spread in any of the environment surveyed. The emerging biological properties of GVL remain elusive. Given that the samples harboring GVL also presented several grapevine viruses and viroids, including *Grapevine rupestris stem pitting-associated virus* (*Foveavirus*), *Grapevine Syrah virus-1* (*Marafivirus*), *Grapevine leafroll-associated virus 3* (*Ampelovirus*), *Grapevine yellow speckle viroid* 1 & 2 (*Apscaviroid*), and *Hop stunt viroid* (*Hostuviroid*), future studies should unravel whether GVL presence is associated to any specific symptoms in grapevine.

**Figure 2.**
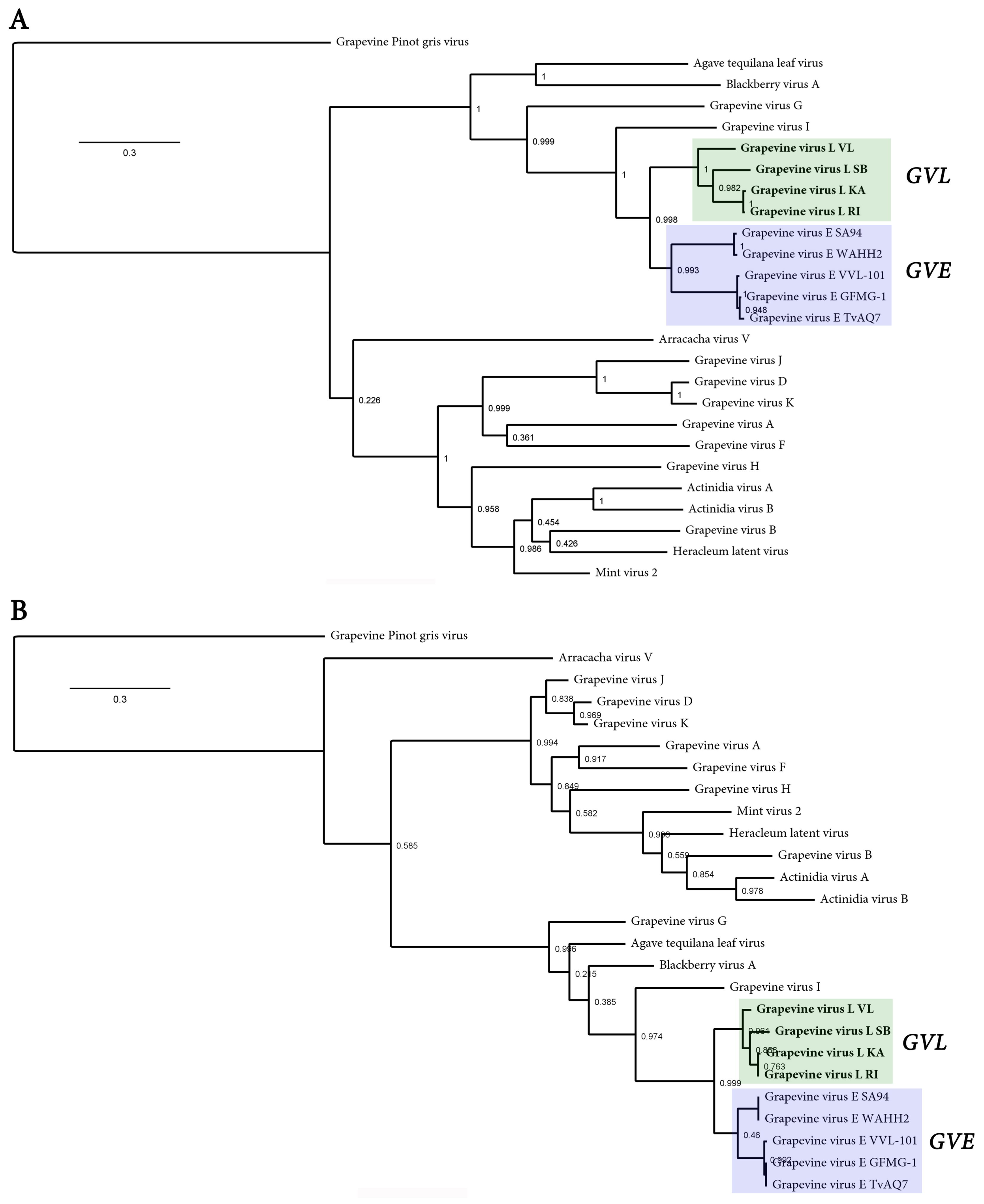
Phylogenetic insights integrating four isolates of grapevine virus L (GVL) and five isolates of grapevine virus E in the context of members of the genus *Vitivirus* based on MAFFT alignments (BLOSUM62 scoring matrix using as best-fit algorithm E-INS-i) of the predicted replicase (**A**) and capsid protein (**B**) followed by maximum likelihood trees generated by FastTree (best-fit model = JTT-Jones-Taylor-Thorton with single rate of evolution for each site = CAT) computing local support values with the Shimodaira-Hasegawa test (SH) and 1,000 resamples). Accession numbers of the corresponding sequences: grapevine virus E (GVE) isolate VVL-101(MF991950), GVE isolate WAHH2 (JX402759), GVE isolate SA94 (GU903012), GVE isolate GFMG-1 (KF588015), GVE isolate TvAQ7 (NC_011106). Additional sequences are indicated in Figure 1.B legend. Pinot gris virus was used as outgroup. Scale bar represents substitutions per site. Node labels are FastTree support values. GVL isolates are depicted in bold.

## Acknowledgements

This work was supported by project PE1131022 of the Instituto Nacional de Tecnologìa Agropecuaria (INTA) and by ANPCyT PICT 2015-1532 and PICT 2016-0429. The funders had no role in study design and analysis, decision to publish, or preparation of the manuscript. The research from New Zealand is funded by New Zealand Winegrowers Inc. and The Ministry of Business, Innovation & Employment. Many thanks to Alfredo Diaz Lara, Department of Plant Pathology, University of California, Davis, CA for his technical assistance.

## Author contributions

HD, SGT and SA designed the study. RSB, DV, RPPA, AGB and MAR, performed experiments, generated and analyzed data. HD integrated the data together with SA. HD wrote the initial draft of the manuscript and DZ, SGT, AGB, RSB, DV, RPPA, MAR and SA revised the manuscript. All authors approved the final version.

## Ethical approval

This article does not contain any studies with human participants or animals performed by any of the authors. Therefore, informed consent was not required for this work.

---

**Supplementary Figure 1.**
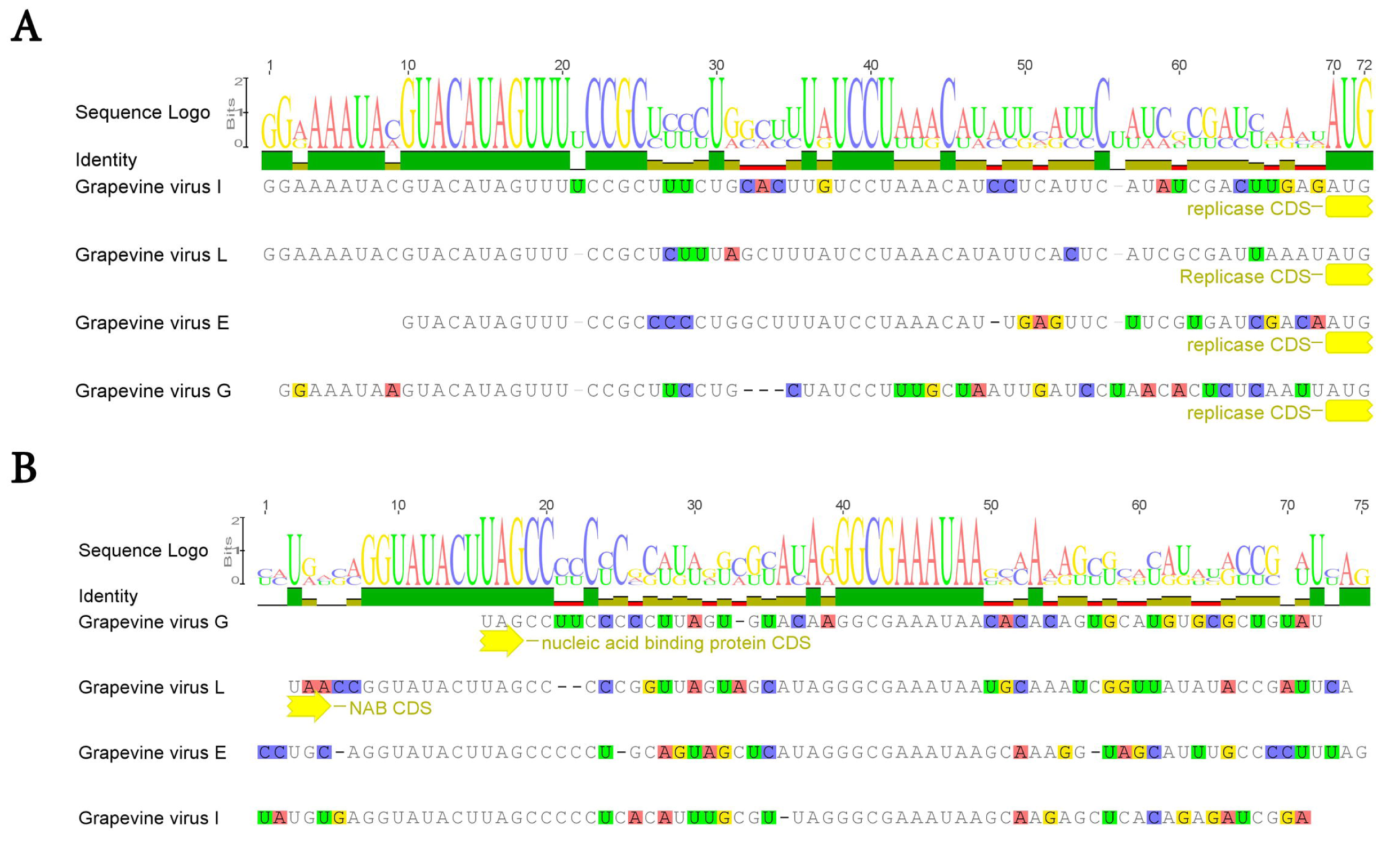
ClustalW alignment of 5’-UTR (**A**) and 3’-UTR (**B**) of GVL, GVE and the proposed GVI and GVG. GVL 5’-UTR terminal 20 nt are identical to that of GVI. At the 3’-UTR in an equidistal position of 3’ terminus a ten nucleotides monomer “GGCGAAAUAA” is conserved among the four virus sequences.

**Supplementary Figure 2.**
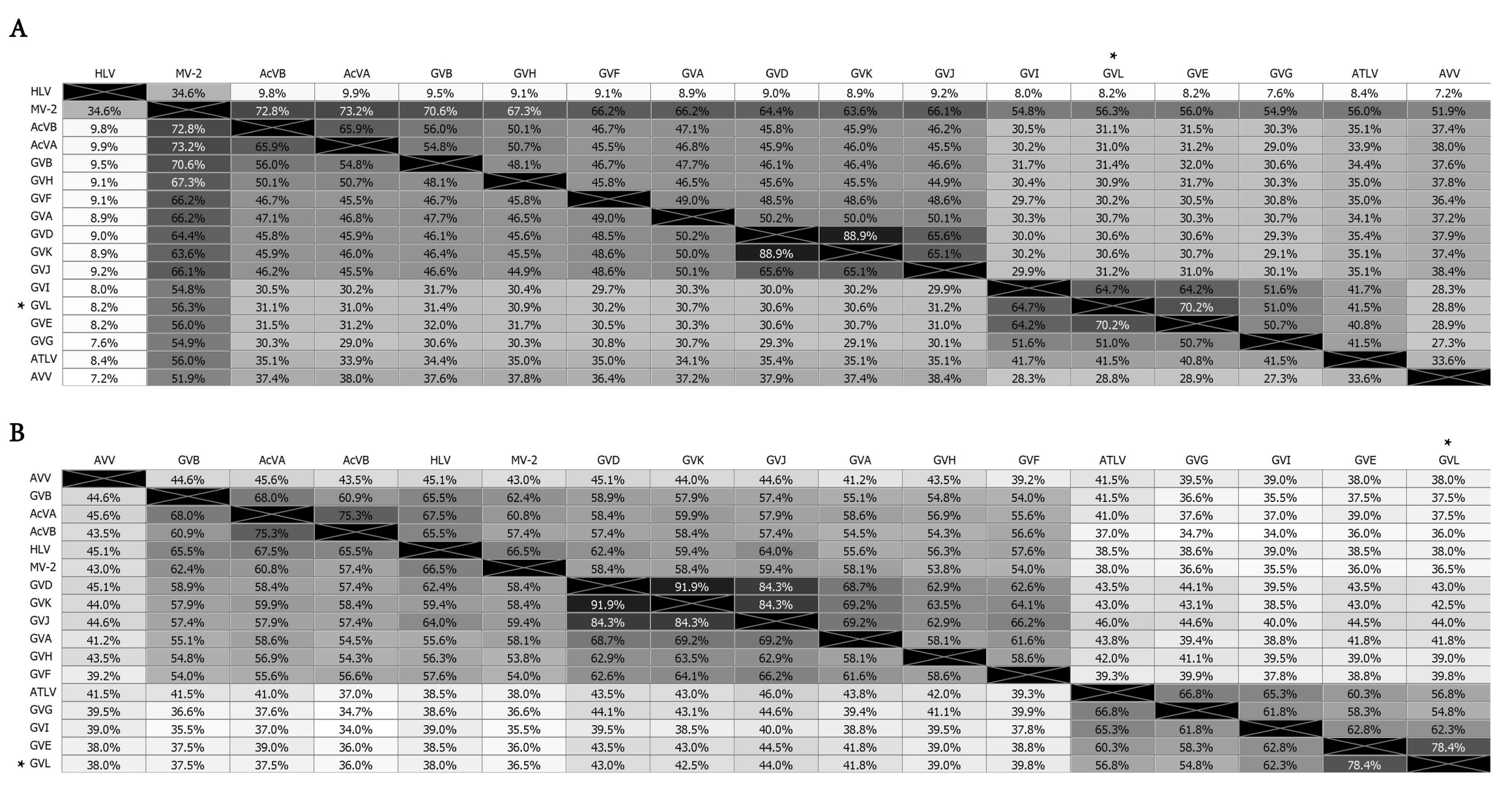
Genetic distances expressed as percentage values and heat-maps of predicted RP (A) and CP (B) of accepted and proposed members of the genus *Vitivirus* and GVL, based on MAFFT alignments (BLOSUM62 scoring matrix using as best-fit algorithm E-INS-I). See Table 1 for abbreviations.

**Supplementary Figure 3.**
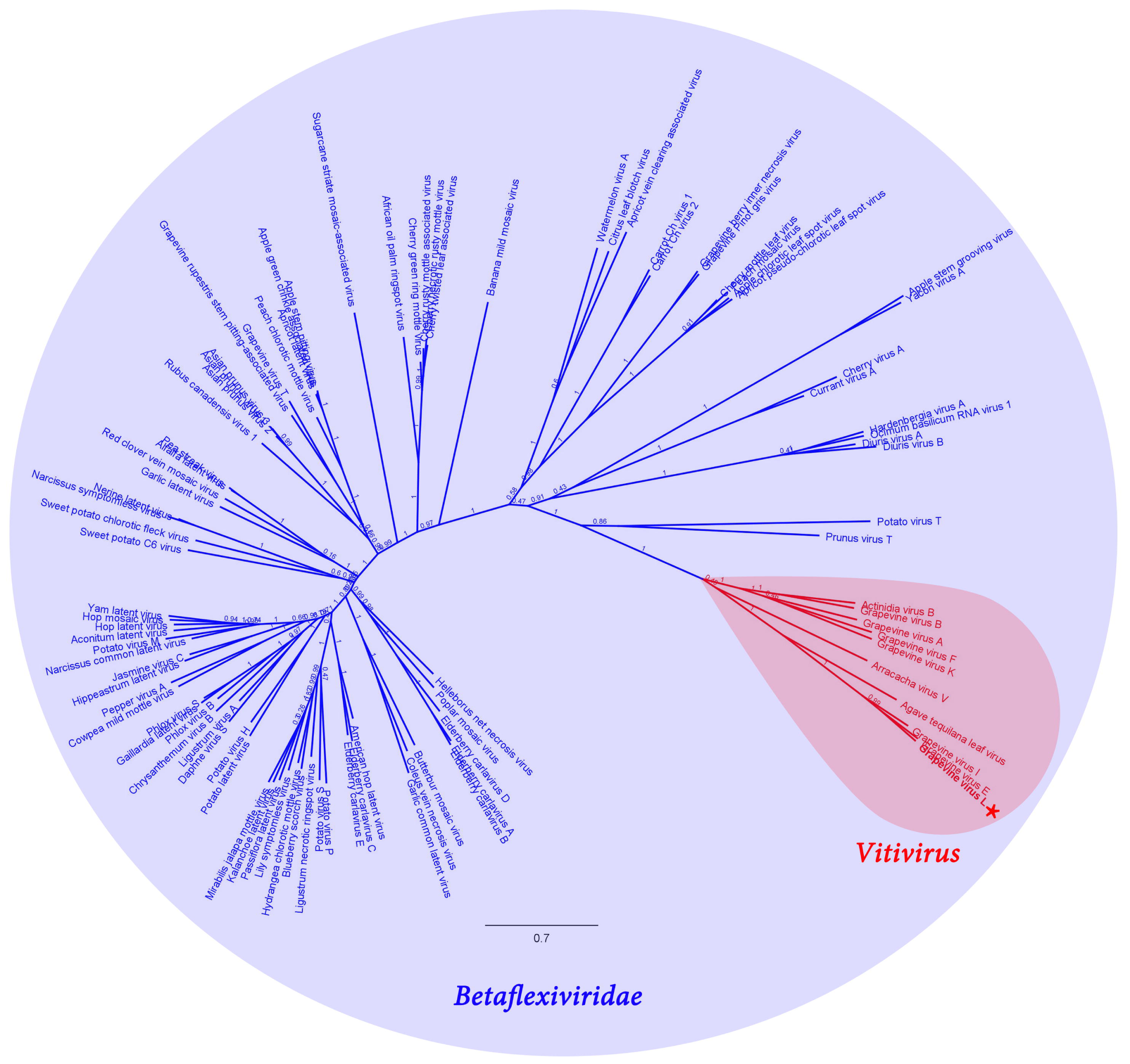
Phylogenetic insights of putative grapevine virus L (GVL) in the context of family *Betaflexiviridae*, based on MAFFT alignments (BLOSUM62 scoring matrix using as best-fit algorithm E-INS-i) of NCBI refseq replicase proteins of *Betaflexiviridae* members, followed by maximum likelihood radial trees generated by FastTree (best-fit model = JTT-Jones-Taylor-Thorton with single rate of evolution for each site = CAT) computing local support values with the Shimodaira-Hasegawa test (SH) and 1,000 resamples). Scale bar represents substitutions per site. *Betaflexiviridae* family is depicted in blue and the monophyletic clade of vitiviruses is shown in red. GVL is indicated with an asterisk.

**Supplementary Figure 4.**
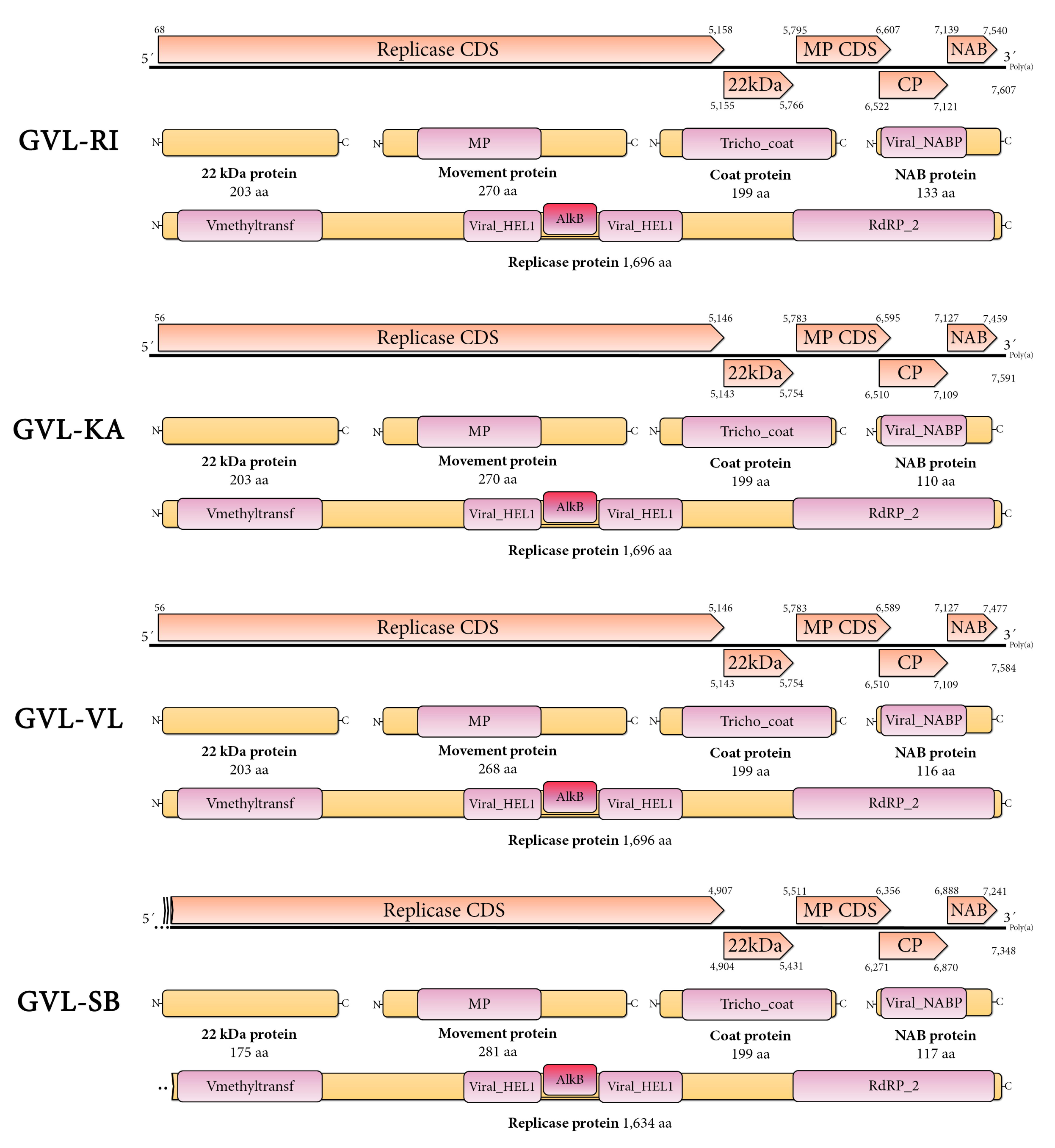
Genomic architecture of the draft genomes of the identified GVL isolates characterized by a monosegmented ssRNA(+) virus encoding for five ORFs arranged as 5’-UTR-RP-HP-MP-CP-NABP-UTR-3’. The predicted product of each ORF is depicted including the associated domains determined by NCBI-CDD. Abbreviations: RP, replicase protein; 22kDa, 22k Dalton hypothetical protein; MP, movement protein; CP, coat protein; NABP, nucleic acid binding protein.

**Supplementary Table 1.** NGS RNA libraries used in this study generated by Chen et al [10]. Stage: sampled grapeberries developmental stage; Total reads: Total raw 50 nt reads generated for each library; GVL virus reads: viral reads detected in each sample; GVL RPM: virus reads per million total library reads.

**Supplementary Table 2.** Variable sites of grapevine virus L identified in the virus RNA reads detected in SRX3144939-SRX3144940 (E-L 35 cv Riesling grapeberries). Polymorphism are indicated in relation with the consensus sequence of the E-L-38 cv Riesling strain determined for SRX3144956. Only variable sites with a frequency >0.5 of virus reads are shown.

**Supplementary Table 3.** Structural highlights and genetic distance of grapevine virus L isolates. Abbreviations: Accession #: NCBI Genbank accession number; GS: Genome size (nt), 5′U: 5′UTR length (nt); RP cds: replicase cds length (nt); HP cds: hypothetical cds length (nt); MP cds: movement protein cds length (nt); CP cds: coat protein cds length (nt); NABP: nucleic acid binding protein cds length (aa); 3′UTR: 3′UTR length excluding the poly A tail (nt). RPd-NABd: genetic distance of genomic region (nt) / predicted gene product (aa) of corresponding isolate with GVL-RI.

